# GlycoEnzOnto: A GlycoEnzyme Pathway and Molecular Function Ontology

**DOI:** 10.1101/2022.06.06.493779

**Authors:** Theodore Groth, Rudiyanto Gunawan, Alexander D. Diehl, Sriram Neelamegham

**Affiliations:** Department of Chemical and Biological Engineering, State University of New York, Buffalo, NY, 14260, USA; Department of Biomedical Informatics, State University of New York, Buffalo, NY, 14260, USA; Department of Biomedical Engineering, State University of New York, Buffalo, NY, 14260, USA; Department of Medicine, University at Buffalo, State University of New York, Buffalo, NY, 14260, USA

**Keywords:** glycoEnzymes, glycosylation pathways, ontology

## Abstract

The ‘glycoEnzymes’ include a set of proteins having related enzymatic, metabolic, transport, structural and cofactor functions. Current there is no established ontology to describe glycoEnzyme properties and to relate them to glycan biosynthesis pathways. We present GlycoEnzOnto, an ontology describing 386 human glycoEnzymes curated along 135 glycosylation pathways, 134 molecular functions and 22 cellular compartments. The pathways described regulate nucleotide-sugar metabolism, glycosyl-substrate/donor transport, glycan biosynthesis, and degradation. The role of each enzyme in the glycosylation initiation, elongation/branching, and capping/termination phases is described. IUPAC linear strings present systematic human/machine readable descriptions of individual reaction steps and enable automated knowledge-based curation of biochemical networks. All GlycoEnzOnto knowledge is integrated with the Gene Ontology (GO) biological processes. GlycoEnzOnto enables improved transcript overrepresentation analyses and glycosylation pathway identification compared to other available schema, e.g. KEGG and Reactome. Overall, GlycoEnzOnto represents a holistic glycoinformatics resource for systems-level analyses.

**Availability:** https://github.com/neel-lab/GlycoEnzOnto

## INTRODUCTION

Glycosylation results in the biosynthesis of complex carbohydrates or glycans on cell-surface and nuclear proteins, and lipids (Neelamegham and Mahal, 2016). During this process, nucleotide-sugar donors (e.g. GDP-fucose) are formed by the metabolic processing of monosaccharides derived from either dietary sugars (commonly glucose and fructose) or from salvage pathways that breakdown recycled glycoconjugates. These nucleotide-sugars are subsequently transported into the endoplasmic reticulum (E.R.) and Golgi where they act as donors for enzymatic reactions catalyzed by a family of enzymes called glycosyltransferases (GTs). These GTs act sequentially to catalyze the transfer of monosaccharides from the nucleotide-sugar donors to target substrates expressed on protein/lipid scaffolds. Such biosynthetic processes result in many types of glycans in humans, with the four major subclasses being: i. N-linked glycans, ii. O-linked glycans (O-GalNAc, O-GlcNAc etc.), iii. glycolipids, and iv. glycosaminoglycans (heparan sulfates, hyaluronan etc.) (Neelamegham *et al*., 2022; Schjoldager *et al*., 2020). The cellular ‘glycome’ encompasses the collection of all glycans.

Systems-based bioinformatics analyses of glycosylation requires formalized knowledge, including ontologies. Among these, GlycoRDF and Glycoconjugate Ontology (GlycoCo) help harmonize glycan structural data from various glycoscience databases (Ranzinger *et al*., 2015; Yamada *et al*., 2021). Here, GlycoRDF describes the framework to model instances of glycogenes and reactions, though individual annotations of glycoEnzymes, pathways and molecular functions are not part of this undertaking. The Glycan Naming and Subsumption Ontology (GNOme) (Edwards, 2021) describes the topological connectivity of monosaccharides and substructures within complex carbohydrates, and this facilitates glycan queries at the GlyGen repository (York *et al*., 2020; Kahsay *et al*., 2020). The genetic Glyco-Disease Ontology (GGDonto) curates information regarding glycoEnzyme dysfunction and related pathways in disease contexts, especially the congenital disorders of glycosylation (Solovieva *et al*., 2018). To date, no existing resource presents a comprehensive glycoEnzyme functional hierarchy.

This manuscript presents GlycoEnzOnto, a manually curated ontology that captures current biological knowledge regarding 386 human GlycoEnzymes and demonstrates its applications in bioinformatics data analyses. These glycoEnzymes are annotated into 135 glycosylation pathway classes, 134 molecular functions and 22 cellular compartments, similar to the organization of the Gene Ontology (GO) (Ashburner *et al*., 2000). In cells, the glycoEnzymes facilitate nucleotide-sugar metabolism, transport, glycan biosynthesis, and degradation. Using biochemical knowledge about their mechanism, individual glycoEnzyme reactions and preferred glycan substrates are manually curated using a novel IUPAC-based string code. In this regard, while previous efforts to describe glycoEnzyme reaction specificity were either XML-based (Liu and Neelamegham, 2014; Liu *et al*., 2013) or LinearCode-based (Krambeck *et al*., 2009; Kellman *et al*., 2020), GlycoEnzOnto uses IUPAC due to its common usage in literature. This both enhances human-readability and systemizes conversion of reaction rules for machine usage. During their curation, the glycoEnzymes were delineated based on their modular contribution to glycosylation: i. ‘initiation’ steps that result in the attachment of the first monosaccharide or oligosaccharide to the protein/lipid, ii. ‘elongation and branching’ reactions which often involve lactosamine chain synthesis and branching, and iii. ‘termination or capping’ steps which prevent further chain extension. Such pathway classifications, when utilized in gene overrepresentation analyses of disease datasets (e.g. from The Cancer Genome Atlas (TCGA) (Grossman *et al*., 2016)), enables the identification of glycan biosynthetic processes that may be dyregulated during disease. Overall, GlycoEnzOnto aims to enable large-scale, high-throughput data analyses in the Glycoscience domain, and enhance sharing of results based on the FAIR (Findable, Accessible, Interoperable and Reusable) principles.

## SYSTEMS AND METHODS

### Instantiating glycoEnzymes in GlycoEnzOnto and integration with Gene Ontology (GO)

386 human glycoEnzymes were curated in GlycoEnzOnto using textbook/handbook references (Taniguchi *et al*., 2014; Varki *et al*., 2017), supplemented with additional knowledge from literature (Neelamegham *et al*., 2022; Schjoldager *et al*., 2020; Huang *et al*., 2021). The curated GlycoEnzOnto.owl is freely available and related data are also presented in table form (**Supplemental Table S1**). In GlycoEnzOnto, UniProt URIs corresponding to proteins and genes are defined under the ‘Protein’ and ‘Gene’ classes. ‘Protein’ instances are linked to ‘Gene’ using the UniProt object property ‘encodedBy’. A symmetric object property ‘encodes’ relates UniProt ‘Protein’ instances to ‘Gene’. Each glycoEnzyme is assigned to corresponding pathway(s) and given a systematic reaction rule string.

Terms in the Gene Ontology (GO) are included in GlycoEnzOnto to integrate existing knowledge regarding glycoEnzyme function alongside pathway, molecular function and compartment data. To this end, GO terms for each glycoEnzyme were obtained from the UniProt SPARQL endpoint. The GO class hierarchies for the glycoEnzymes associated with biological processes, molecular functions and cellular components were extracted using the MIREOT method in ROBOT (Jackson *et al*., 2019), and stored in a ‘glycoenzyme GO term subset file’. Relevant Relation Ontology (RO) terms were similarly extracted into a ‘Relation Ontology subset file’. Next, the Basic Formal Ontology (BFO), ‘Relation Ontology subset’, and ‘glycoenzyme GO term subset’ files were imported into GlycoEnzOnto. Three manual steps were undertaken to integrate knowledge and establish GlycoEnzOnto-GO relations: i) The top-level GlycoEnzOnto class ‘glycosylation-related pathway’ was made a subclass of GO ‘biological process’. ii) UniProt Protein and Gene classes were made ‘material entity’ instances from the BFO. iii) The GlycoEnzOnto classes were manually associated with GO biological processes by creating equivalence axioms that utilized primarily two object properties from Relation Ontology: the ‘occurrent part of’ (RO:0002012) and the ‘encompasses’ object property (RO:0002085). In this last step, we ensured the existence of close relationships between the GO term description and GlycoEnzyOnto pathway. Whereas all enzymes in GlycoEnzyOnto pathway were a subset of the GO term in the case of ‘occurrent part of’ relation, the exact opposite was the case for the ‘encompasses’ relation. Following the above, 58 ‘occurrent part of’ and two ‘encompasses’ relations were established. Overall, linking GlycoEnzOnto and GO terms using the RO contextualizes glycan structure biosynthetic pathways with biological process knowledge.

### GlycoEnzyme reaction rule strings

A concise IUPAC-based, human-readable, glycoEnzyme reaction rule language was developed in order to describe carbohydrate transformation(s) based on existing biological knowledge. Such reaction rules enabled overlaying of gene expression data on glycosylation knowledge for the purpose of pathway/network enrichment analysis. Here, reaction rules were instantiated as “reaction rule” (genzo:0262), and constraints as “reaction constraint” (genzo:0257).

Five ‘reaction types’ were described in the reaction rule strings, with curly braces used to enclose monosaccharides or substituents that are being transformed (**Table 1**). Such transformations are implicitly assumed to occur at the non-reducing terminus of substrates. In ‘addition’ reactions, monosaccharides placed in curly braces are added to the remaining substrate. In ‘subtraction’ reactions, the entities in curly brackets are preceded by ‘!’ (exclamation mark) to indicate deletion. A double-headed arrow ‘<->’ indicates reaction reversibility, whereas the unidirectional arrow symbol ‘->’ indicates ‘substitution’ which would occur during isomerization or other reactions where one reactant is replaced by another. Finally, ‘transport’ from one compartment to another is described by enclosing the source and destination compartment names within brackets.

**Table 1.**
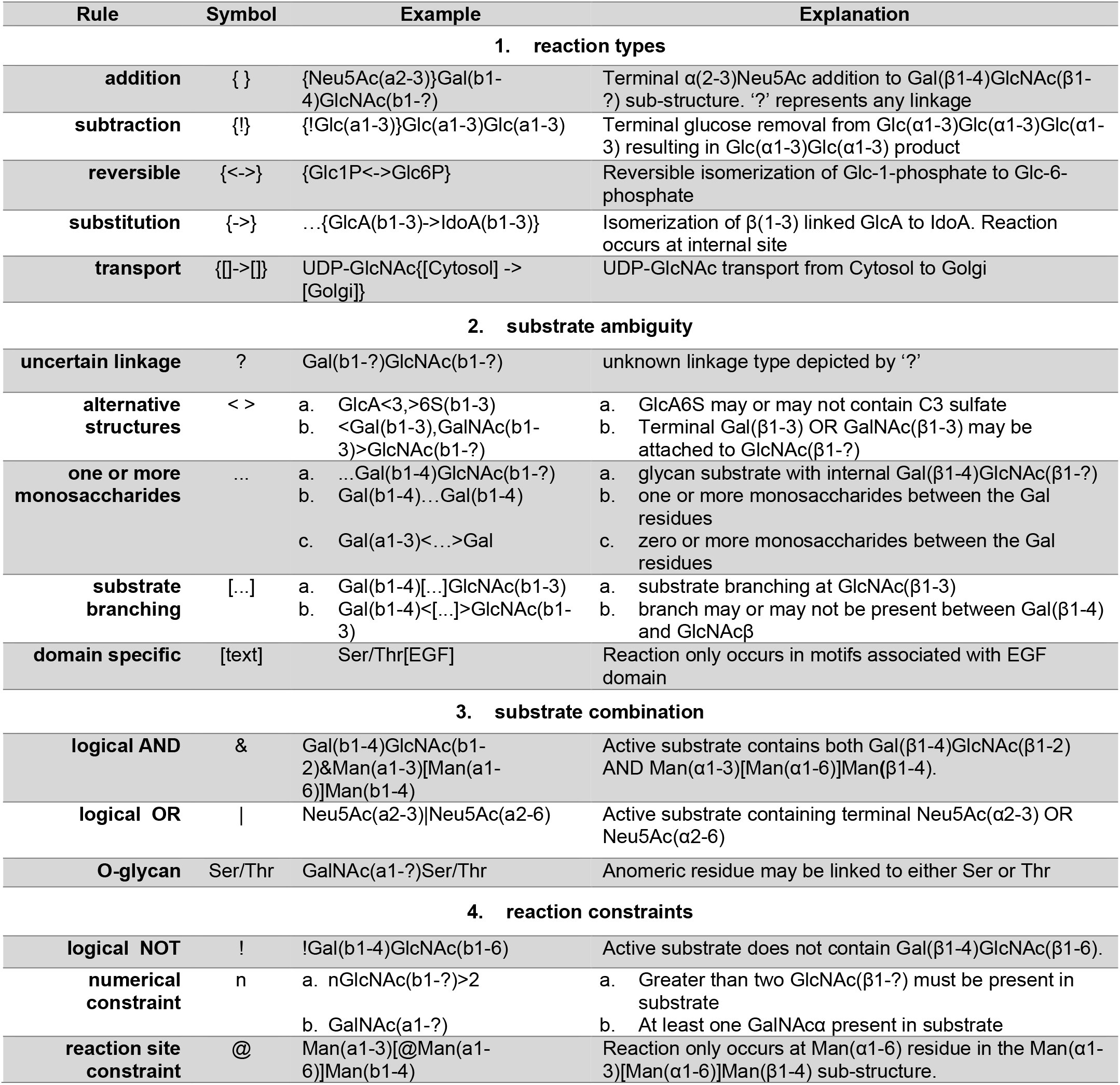
Symbols and constraints to depict glycoEnzyme substrate specificity and reactions.

GlycoEnzymes exhibit group-specific, necessitating the need to describe ‘substrate ambiguity’ to encode structurally similar/equivalent entities. To describe this, ‘?’ is used to depict uncertain substrate bond linkages. Angle brackets (‘<>‘) enclose one or more sub-structures that may or may not be part of a given substrate. The continuation symbol (‘…’) indicates the presence of one monosaccharide or arbitrary glycan sub-structure. If present at the non-reducing terminal, this is used to indicate an internal reaction site. When included in the middle of the substrate, this is to shorten the structure presentation in order to reduce text and focus on more critical features. Examples of continuity operator usage are presented in Table 1, including examples where they are combined with angle bracket (‘<…>’) to indicate zero or more sub-structures. Similarly, the continuation symbol in square brackets (‘[…]’) indicates the presence of obligatory branch(s) in the substrate. Reactions occurring in specific protein domains or motifs are included in these reaction descriptors within square text brackets.

‘Substrate combination’ using logical operations enables more complex substrate definitions. For example, the logical ‘AND’ or ‘&’ symbol is used to indicate the presence of multiple, critical epitopes in a single substrate. The presence of multiple reactive substrates/epitopes is described using the logical ‘OR’ (symbolically shown as ‘|’). Here the presence of either groups would make the substrate reactive.

‘Reaction constraints’ limit specific reaction types. Here, the logical ‘NOT’ or ‘!’ operator specifies glycan structure or sub-structures that prevent biochemical transformation. The number of allowed repeating structures/monosaccharides in a reactive substrate is defined using the symbol ‘n’. The ‘@’ operand is used to restrict the formation of specific products, and this is often used in conjunction with the logical NOT operator. For instance, β1-4 galactosyltransferases (β4GalTs) can act on any terminal GlcNAc, except for bisecting GlcNAc of N-linked glycans. The constraint ‘!@GlcNAc(b1-4)[…]Man(b1-4)’ is used to describe this property. Overall, Table 1 aims to simplify more complicated substrate and product rules described elsewhere (Liu and Neelamegham, 2014; Kellman *et al*., 2020). Codes used to parse reaction rules and constraints are available at the GlycoEnzOnto Github repository.

### Overrepresentation analyses

The ability of GlycoEnzOnto to distill differential expression in an overrepresentation analysis was compared to Reactome, GO, and KEGG glycosylation pathway descriptions (Ashburner *et al*., 2000; Kanehisa and Goto, 2000; Jassal *et al*., 2020). To this end, we extracted analogous pathways and biological processes from these three resources. The detailed gene sets and data collection method used for each repository is fully described in **Supplemental Table S2A-S2C**.

Differential expression analysis was performed on the TCGA breast invasive carcinoma dataset, where changes in gene expression in 81 Her2^+^ breast cancer patients was compared to 113 normal tissue samples. The differential expression analysis was performed by fitting a linear mixed model in the DREAM package (Hoffman and Roussos, 2021), where fixed effects were the PAM50 molecular subtypes of breast cancer and the tumor sample purity (Yoshihara *et al*., 2013), and the random effects were age and the source of the biological material. Enrichments were performed using the entire human transcriptome (22,686 genes) as the universe and the glycoEnzymes (386 genes) as the universe. Differentially expressed genes were defined as having an absolute log-fold-change > log(2), and an adjusted-p-value ≤ 0.05. Finally, overrepresentation analyses were performed to identify which glycosylation pathways were up-/down-regulated in Her2^+^ breast cancer with respect to normal tissue. This was accomplished using Fisher’s exact test, where the ratio of dysregulated glycogenes within a pathway was compared to background dysregulation rates of the entire transcriptome, as well as the glycogene transcriptome.

## IMPLEMENTATION

### GlycoEnzyme instances in GlycoEnzOnto

A total of 386 glycoEnzymes were manually annotated in GlycoEnzOnto and most were given a corresponding reaction rule string. GlycoEnzOnto was cross-referenced with the Gene Ontology (GO) and UniProt knowledgebases in order to contextualize glycosylation knowledge in a larger biological framework. This is illustrated using the (α1-3)fucosyltransferase FUT4 as an example (**Figure 1**). Here, the accession number for FUT4 (‘P22803’) is instantiated under the ‘Protein’ class, which is defined in the UniProt ontological namespace. The inverse object properties ‘encodedBy’ and ‘encodes’ relate UniProt proteins to glycogene instances and vice versa. FUT4 is annotated to GlycoEnzOnto pathways (yellow), as well as GO biological process, molecular function, and cellular compartment terms (green). GlycoEnzOnto pathways which take place as part of GO biological processes were linked to one another through the ‘occurrent part of’ object property relation. FUT4, for instance, is annotated to the GlycoEnzOnto “terminal fucosylation biosynthetic pathway” (genzo:0063), which is related to the congruous GO biological process “fucosylation” (GO:00036065) through the ‘occurrent part of’ relationship. The molecular function class, “4-galactosyl-N-acetylglucosaminide-3-alpha-L-fucosyltransferase activity” (GO:0017083) and “trans-Golgi network” cellular compartment (GO:0005802) were also annotated to FUT4. In terms of ‘reaction rules’, FUT4 fucosylates a variety of glycans preferentially on GlcNAcβ in internal type-II LacNAc structures, and it has a ‘capping’ role. However, it does not act proximal to I-branching locations as illustrated in the ‘constraint’ class. The GlycoEnzOnto object properties ‘has reaction rule’ and ‘has constraint’ are used to encode the reaction rule and constraints for FUT4. The instantiation of glycoEnzymes as UniProt ‘Protein’ and ‘Gene’ URIs, along with their annotations to various aspects of the GO makes it easy to use GlycoEnzOnto as a resource to obtain cross-reference data such as EC number, peptide sequence, literature sources, etc. (Uniprot Consortium, 2018).

**Figure 1.**
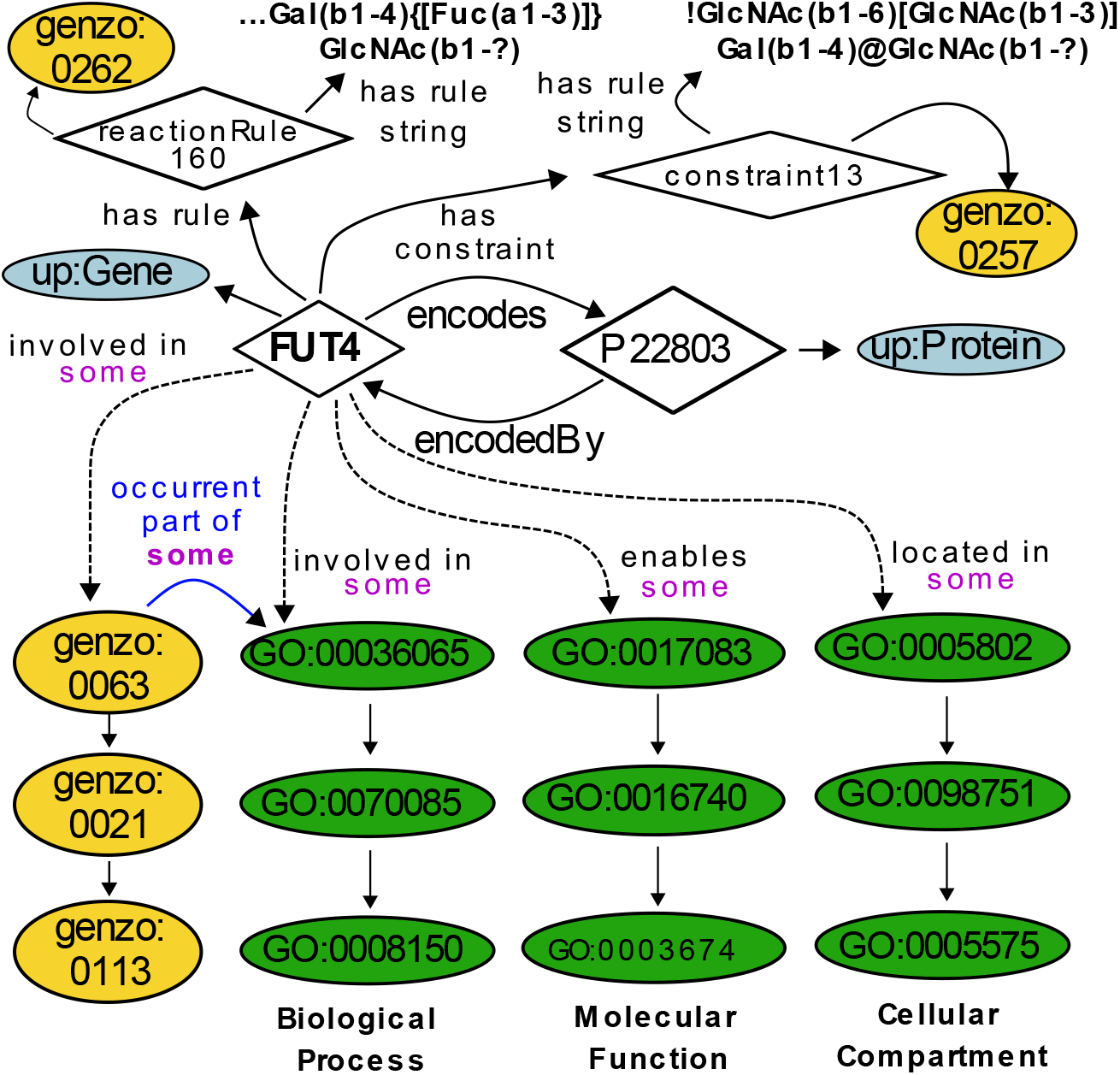
FUT4 description in GlycoEnzOnto. Classes are depicted as ellipses and instances as diamonds. Yellow ellipses represent classes contributed by GlycoEnzOnto, light blue represents the UniProt namespace, and green represents the gene ontology (GO) namespace. Unless labeled, all edges connecting classes and instances have “rdfs:subClassOf” relationship. Arrows depict the object properties linking individuals and classes.

### GlycoEnzOnto molecular functions, pathways and compartments

GlycoEnzOnto presents the classification of 386 glycoEnzymes based on molecular function, pathway and compartment (**Figure 2**). **Fig. 2A** illustrates the glycoEnzyme molecular function hierarchy. This describes the reaction types and other processes mediated by GlycoEnzOnto proteins. This includes six groups: i. ‘transferase’, which either transfer monosaccharides to glycans (‘glycosyltransferases’) or various chemical substituents to monosaccharides (‘other transferases’); ii. ‘modifying enzyme’, that either alter the stereochemistry of substituents or otherwise modify the chemical substituents attached to monosaccharides; iii. ‘glycosidase’, which hydrolyze monosaccharides, often in the lysosomal compartment to contribute to the salvage pathways; iv. ‘transporters’ that shuttle nucleotide-sugars from the cytoplasm to the E.R. or Golgi to provide donors for glycosylation. They also facilitate the transport of the sulfate donor PAPS and monosaccharides from different compartments; v. ‘regulator’, which have non-catalytic functions, but which interact with other GlycoEnzOnto entries to modulate glycosylation; and vi. ‘putative or inactive’ glycoEnzymes whose function is yet to be fully elucidated.

**Figure 2.**
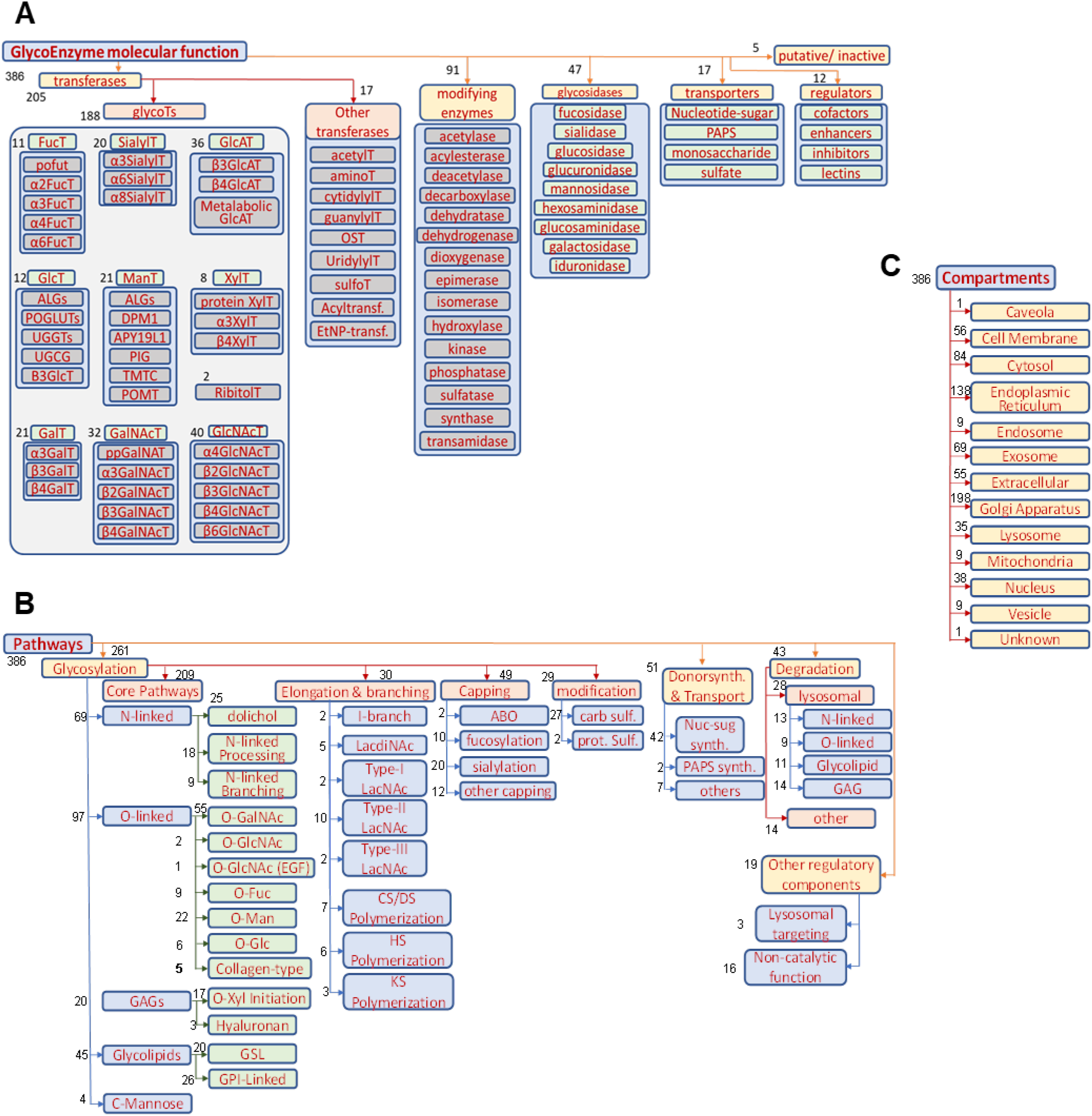
GlycoEnzOnto Classification. A total of 386 glycoEnzymes were classified based on either molecular function (panel **A**), pathway (panel **B**) or compartment (panel **C**). The number of members in each (sub-)group is shown using black text next to the individual entries. Some members may appear in more than one subgroup.

**Fig. 2B** illustrates the biological pathway-based classification of the glycoEnzymes. A majority of the enzymes in this hierarchy facilitate glycosylation, while others mediate sugar-nucleotide biosynthesis and transport, degradation or have other regulatory roles. Here, many of the glycoEnzymes in GlycoEnzOnto ‘initiate’ the biosynthesis of major mammalian glycan families: i. N-linked glycans, ii. O-linked glycans, iii. Glycosaminoglycans (GAGs) and iv. Glycolipids. The O-glycosylation pathways are further broken into common O-GalNAc and O-GlcNAc type modifications, and rarer carbohydrate modifications associated with epidermal growth factor-like repeat domains (EOGT, POFUT1 mediated O-Fuc, O-Glc), Thrombospondin type 1 repeats (POFUT2 mediated O-Fuc), Collagen-type, Cadherin-associated (TMTC-type), and other O-Mannose modifications. The GAGs include the four principal families of hyaluronans, keratan sulfate, heparan sulfate and chondroitin/dermatan sulfate. Also, included are the pathways regulating GPI-anchored protein biosynthesis and C-Mannosylation.

The glycan precursor formed during the ‘initiation’ step may be further modified during ‘elongation and branching’. This results in increased carbohydrate size. The subclasses in this group include the genes regulating the biosynthesis of: i. ‘Type-I/-II/-III LacNAc’ structures, ii. ‘LacDiNAc’ chains, iii. (β1-6)GclNAc branches, and iv. glycosaminoglycan polymerization (for chondroitin/dermatan, heparan and keratin sulfates). In the final ‘capping’ step, the glycoEnzymes generate modifications that preclude further chain extension, commonly using the sialyltransferase and fucosyltransferase enzymes. This often results in the biosynthesis of named carbohydrate epitopes, such as sialyl Lewis-X, blood group antigens and HNK-1.

‘Compartments’ are curated for each of the GlycoEnzymes (**Figure 2C**). Supporting data come from Gene Ontology, UniProt, and other references. This classification includes 22 primary compartments and additional sub-compartments in the case of Golgi, when such data are available. Delineating the glycoEnzymes into compartments, provides context for the glycosylation reactions and enables grouping during modeling tasks.

### Glycosylation pathways generated using glycoEnzyme reaction rules

The GlycoEnzOnto reaction rules enable *in silico* glycosylation pathway generation for various glycoconjugate types (**Figure 3**). **Fig. 3A** illustrates various reaction types accommodated by GlycoEnzOnto: addition, subtraction, transport etc. Using python scripts that processes input glycans, reaction rules and constraint strings defined in GlycoEnzOnto, it is possible to generate glycan biosynthesis pathways like the N-glycosylation pathway illustrated in **Fig. 3B**. The inputs for this example include a set of eight complex N-linked glycans (shown within box) and five glycoEnzymes (MGAT2, MGAT4A/B/C, B4GALT1, ST6GAL1, and ST3GAL1). The script processes each of the input glycans to infer products described as IUPAC condensed outputs. While one-cycle of substrate production is presented for the sake of illustration, the product inference operation may be repeated to generate larger networks. Such network generation may include stopping conditions, e.g. specification of maximum allowable glycan mass, or final glycan structure(s). Overall, the glycoEnzyme definitions of GlycoEnzOnto allow efficiency reaction network generation.

**Figure 3.**
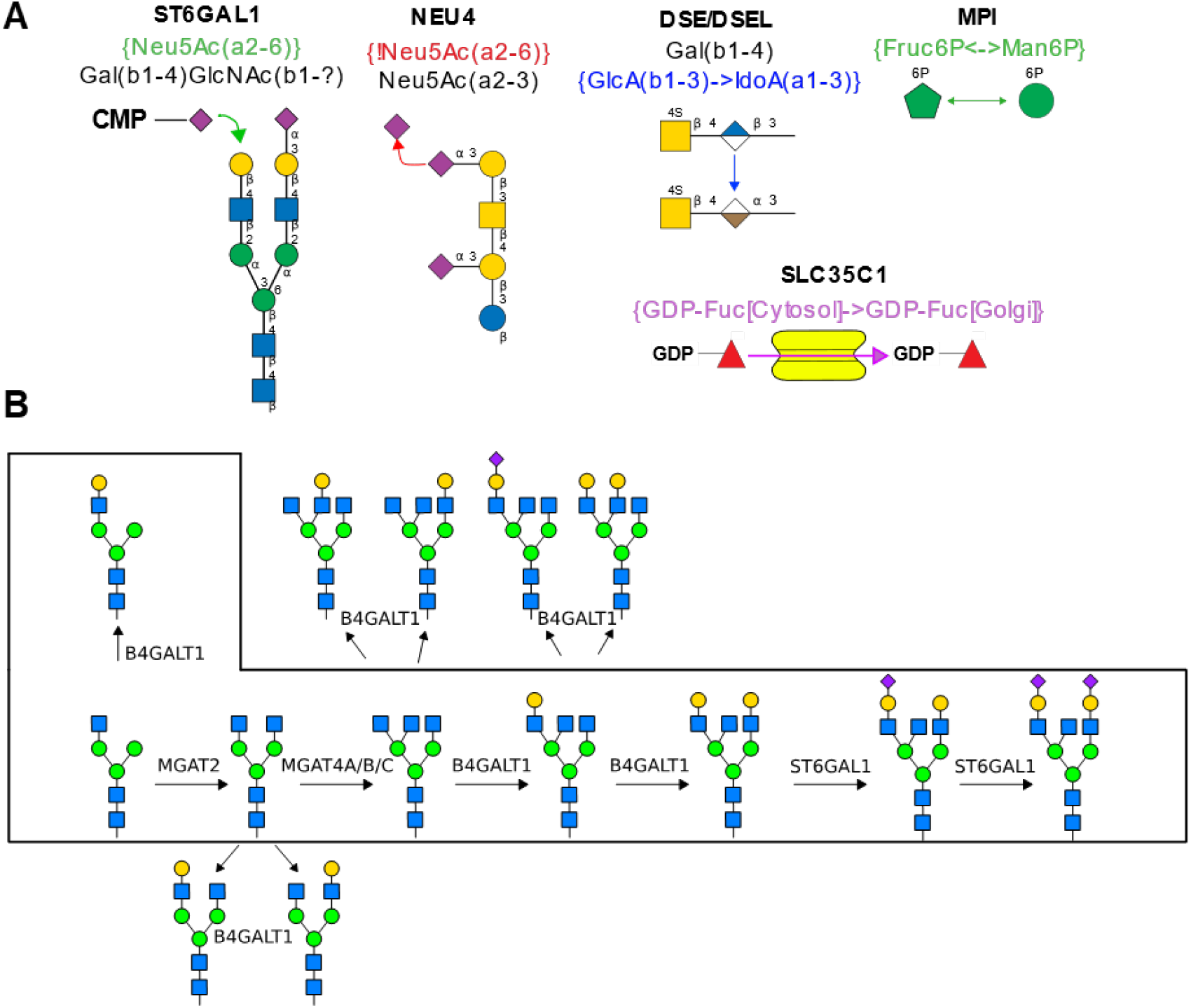
N-linked glycosylation pathway generation using GlycoEnzOnto reaction rules and constraints. **A.** Depiction of five types of reactions described by GlycoEnzOnto (see Table 1 for more details). These are presented using specific glycoEnzymes examples: ST6Gal1, Neu4, DSE/DSEL, MPI and SLC35C1 (all part of GlycoEnzOnto). **B**. Glycan structures shown in box were seeded into the network generation algorithm, along with enzymes B4GALT1, MGAT2, MGAT4, ST6GAL1 and ST3GAL1. Results from first cycle of product inference is shown, with newly generated glycans outside of the boxed area. Note that, in this example, the (α2-3)sialytransferase ST3GAL1 does not process any of the input glycans as they do not contain the required reactive Type III lactosamine substrate. Thus, it is not part of the figure. Additional cycles may be performed to generate a larger network.

### GlycoEnzOnto enables enhanced enrichment analysis

We used gene expression data to evaluate the ability of GlycoEnzOnto to discover glycan pathways dysregulated during disease. Results were compared with the same analysis performed using GO, KEGG and Reactome (**Figure 4**). The analysis focused on the TCGA transcriptome profile of Her2+ breast cancer tissue with respect to normal breast tissue. Such analysis was performed both upon considering all human transcripts (Fig. 4A-B), and when considering only the glycogenes curated in GlycoEnzOnto (Fig. 4C-D).

**Figure 4.**
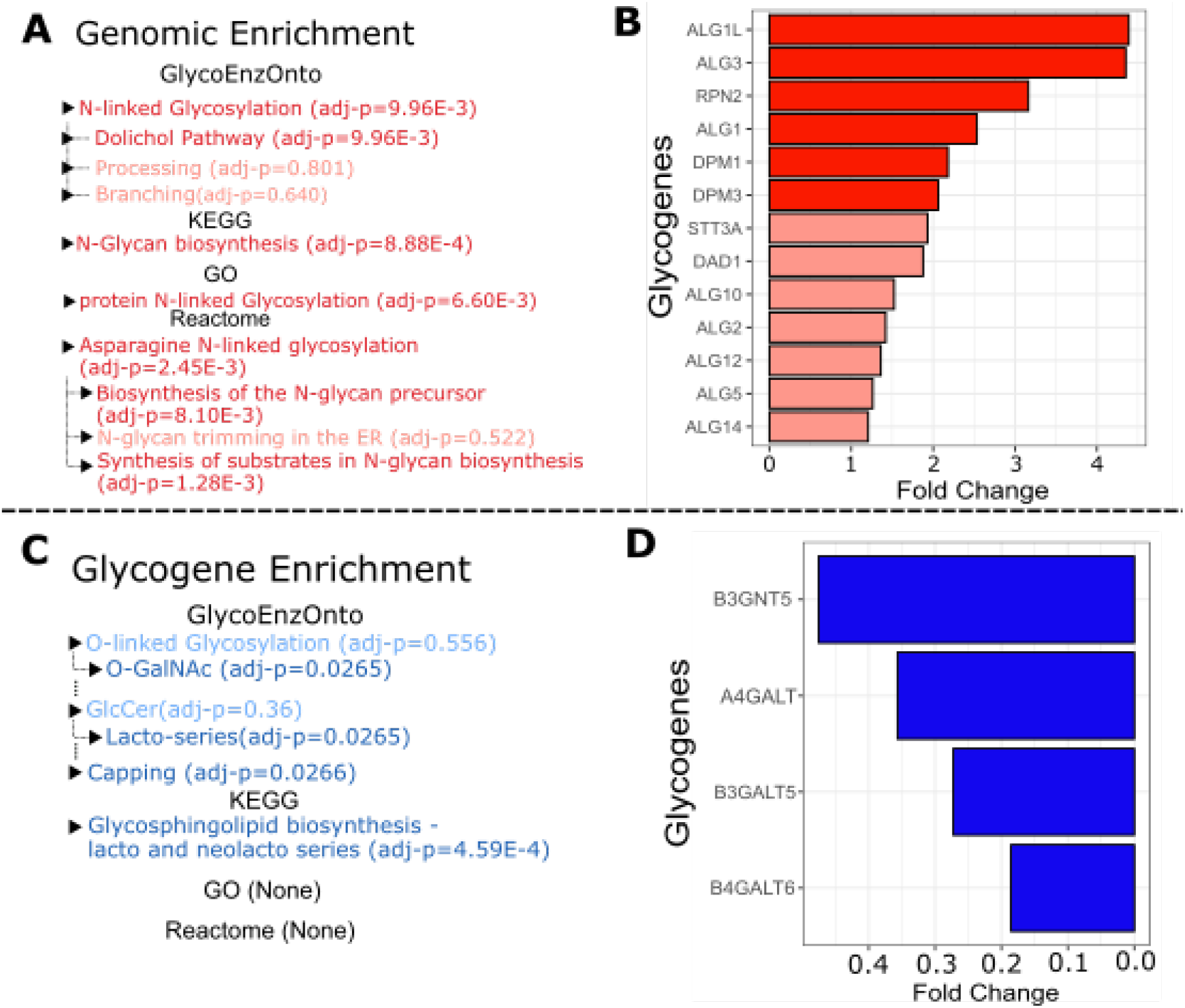
Glycosylation pathway enrichment. Enrichment tests were performed to compare the transcriptome of normal breast tissue with Her2^+^ cancer tissue. Differential expression analysis was performed upon both considering the entire human transcriptome (26,686 genes) as the universe (panels **A, B**), and upon just considering the 386 glycoEnzymes (panels **C, D**). Upregulated and downregulated pathways in these panels are highlighted in bright red and blue, respectively (panel **A, C**). Lighter shades indicate pathways that are altered, but not statistically significantly modified. Bar plots illustrate increased glycogenes that are part of the dolichol biosynthesis pathway in Her2^+^ breast tissue (panel **B**), and lacto-series initiation enzymes that are decreased during cancer (panel **D**).

Upon considering all human transcripts, all four resources detected that N-linked glycosylation was disproportionately upregulated in Her2^+^ tissue (Fig. 4A). GlycoEnzOnto and Reactome, more specifically, detected that the dolichol precursor pathway was dysregulated, whereas this level of detailed inference was not possible using GO and KEGG due to incomplete curation of sub-pathways. While the Reactome ‘Biosynthesis of the N-glycan precursor’ and ‘Synthesis of substrates in N-glycan biosynthesis’ pathways were upregulated, only a small fraction of the genes in these classifications are actually involved in dolichol-precursor biosynthesis. Most of the genes in these classifications were out-of-scope as they involved metabolic processing, nucleotide-sugar biosynthesis (GDP-fucose, CMP-sialic acid etc.), sialylation and sialidase activity (Supplemental Table S2C). In agreement with GlycoEnzOnto over-expression analysis, indeed, all thirteen genes regulating N-linked dolichol precursor synthesis were upregulated in Her2+ tissue with six of them being statistically significant (Fig. 4B). Significantly upregulated genes include several mannosyltransferases (ALG1, ALG1L, ALG3), Man-Dol-P donor synthesis enzymes (DPM1, DPM3) and RPN2 which regulates dolichol precursor transfer onto asparagine on the nascent protein. As the dolichol precursor pathway proceeds in a linear fashion, the overrepresentation analysis suggests that dolichol biosynthesis may be increased during Her2^+^ breast cancer. Consistent with this, glycomics profiling shows elevated high mannose glycans in breast tumors and cancer cells (Tan *et al*., 2014; Li *et al*., 2019). Here, the increased flux of N-linked glycan biosynthesis may result in the presence of untrimmed, disease-associated oligomannose structures.

When using the glycoEnzymes as the universe for enrichment tests instead of the entire transcriptome, several GlycoEnzOnto pathways were found to be downregulated in Her2+ breast cancer including: i) lactosylceramide synthesis, ii) O-GalNAc glycosylation, and iii) glycan capping (Fig. 4C). KEGG analysis also showed downregulation of lacto and neolacto-series glycans during breast cancer. However, the glycolipid biosynthesis enzymes in KEGG included entities regulating glycolipid initiation, branching and extension, along with several capping fucosyl- and sialyl-transferases. In contrast to this ambiguity, the GlycoEnzOnto ontology suggests that the impact of cancer is likely to be more severe on lactosyl-ceramide biosynthesis initiating enzymes: A4GALT5, B3GALT5, B3GNT5, and B4GALT6 (Fig 4D). With respect to the O-GalNAc pathway, we observed alterations in several related glycogenes including the downregulation of GCNT4, GALNT13, GALNT16 and C1GALT1, and upregulation of ST3GAL1, ST6GALNAC4 and GAL3ST2 in Her2^+^ breast cancer tissue. These changes would promote the formation of sialyl-T (Neu5Ac(α2-3)Gal(β1-3)GalNAcα) and sialyl-Tn (Neu5Ac(α2-6)GalNAcα) type glycans that have been reported to be augmented during breast cancer (Burchell et al., 2018; Patil et al., 2014). In addition to the above, the expression of several sialyltransferases (ST8Sia-1,-2,-6, ST3Gal-6) and fucosyltransferases (FUT-1, -4, -9, -10) were dysregulated in Her2+ tissue. In contrast to GlycoEnzOnto, no significant glycosylation pathway alterations could be inferred upon using Reactome or GO knowledge. Overall, the pathway-based description of glycoEnzymes in GlycoEnzOnto enabled a nuanced analysis of glycoEnzymes that participate in the synthesis of various cancer glycoconjugates.

## DISCUSSION

Several resources curate glycosylation related protein functions and pathways including Reactome (Jassal *et al*., 2020), KEGG (Kanehisa and Goto, 2000), GO (Ashburner *et al*., 2000) and UniProt (Uniprot Consortium, 2018). Additional databases like Rhea use ontology terms to curate glycoenzyme reactions using ChEBI molecular identifiers as glycan substrates (Bansal *et al*., 2021). While valuable, these resources currently lack sufficient glycosylation biosynthetic knowledge. In particular, they do not capture the hierarchical nature of glycosylation which includes initiation, elongation/branching, and capping/termination steps. GlycoEnzOnto addresses this gap. While focused on human biology, much of the knowledge and data analysis framework may also be applied to other mammals. For example, this same ontology could be applied to mice after incorporating a few enzymes that are missing in humans like CMAH (Cytidine monophospho-N-acetylneuraminic acid hydroxylase) and A3GALT2 (Alpha-1,3-galactosyltransferase 2).

GlycoEnzOnto curates reaction rules for 386 glycoEnzymes using linear strings that are derived from IUPAC-condensed nomenclature. This schema includes three critical components that describe: i) enzyme specific substrates and related substrate ambiguity; ii) five types of biochemical reactions relevant to glycobiology; and iii) reaction constraints that limits the nature of substrate transformation. The ability to parse the rules and constraints efficiently to generate a glycosylation reaction network was demonstrated for the case of N-linked glycosylation. The scope of this endeavor may be extended to other glycoconjugate types. The ability to automate the generation of glycosylation reaction networks allows the simulation of biochemical reaction networks and fitting with glycan structure experimental data (Krambeck *et al*., 2009; Liu *et al*., 2008; Spahn *et al*., 2016). It allows the comprehensive linking of biochemical pathways with metabolic processes (Hutter *et al*., 2018). Transcriptomic data can also be superimposed on such networks in order to derive glycogene-glycan expression relationships (Huang *et al*., 2021).

The new ontology enables overexpression analysis by taking advantage of the classification of the glycoEnzymes into various molecular functions and biological processes. We illustrate this using a breast cancer example that helps discover a broad range of potential cancer-associated dysregulated pathways. During such analysis, the GlycoEnzOnto pathway hierarchy enabled more nuanced identification of dysregulated pathways compared to GO, KEGG and Reactome. While a single test case is presented here for the purpose of illustration, a larger analysis of all 33 cancer types reported in TCGA and their relation to glycan dysregulation will follow shortly. Thus, the proposed method is designed to generate precise glycoscience hypotheses using gene expression data that may be tested in a wet-lab setting. It also aims to use mathematical analyses to uncover glycoEnzyme relationships within cells that cannot be readily measured *in vivo*.

The availability of GlycoEnzyme biochemical knowledge in OWL format using GlycoEnzOnto can allow the integration of glycoEnzyme knowledge into glycan structure databases, thus facilitating the development of glycan reaction networks stored in RDF format based on the provided reaction rules. Collaborative efforts to integrate GlycoEnzOnto terminology into the gene ontology hierarchy, and also present these data at the GlyGen resource. Overall, the standardization of glycosylation reaction rules, the development of related pathway maps and the usage of this framework for glycan enrichment analysis represents the starting point for the development of systems level knowledge of glycosylation processes.

## Acknowledgement

We thank Dr. Yusen Zhou (University of Pennsylvania) for initiating elements of this project.

